# In silico multi-epitope vaccine against covid19 showing effective interaction with HLA-B*15:03

**DOI:** 10.1101/2020.06.10.143545

**Authors:** Muniba Faiza, Tariq Abdullah, Jose Franklin Calderon-Tantalean, Manish Ravindranath Upadhyay, Abdelrahman H. Abdelmoneim, Fareeha Akram, Bhupender Singh Thakur, Ibrahim Abdulaziz, Chimaobi James Ononamadu, Dina Abdelazim Ghoraba, Saba Munawar, MD Fakhrul Islam Faruque, Collins Kigen, Abhishek Sharma, Ashwani Kumar, Aqsa Khalid, Ali Gharip, Ankit Gupta, Manne Manikumar, Uma Chaudhary

**Affiliations:** Bioinformatics Review, IQL Technologies Pvt. Ltd., New Delhi, 110025, India; Editor, Bioinformatics Review, IQL Technologies Pvt. Ltd., New Delhi, 110025, India; Department of parasitology, Institute of biomedical sciences, University of São Paulo, São Paulo, Brazil; M.Sc. Bioinformatics, Department of Bioinformatics, G.N.I.R.D. Guru Nanak Khalsa College Nathalal Parekh Marg, Matunga East, Mumbai, Maharashtra, India; M.D. Clinical Immunology resident, Sudan Medical Specialization Board, Sudan; BS Bioinformatics, Department of Computer Science, University of Agriculture, Faisalabad, Pakistan; M.Sc. School of Bioengineering and Biosciences, Lovely professional University, Punjab, India; B. Tech. Biochemistry, School of Life Science, Modibbo Adama University of Technology, Yola, Nigeria; Lecturer, Department of Biochemistry and Forensic Science, Nigeria Police Academy Wudil, PMB 3474, Kano State, Nigeria; Department of Neurosurgery, Kasr Alainy Medical School Teaching Hospitals, Faculty of Medicine and University Hospitals, Cairo University, Cairo, Egypt; Research Centre for Modeling and Simulations, National University of Science and Technology Pakistan; Bioinformatics Review-nCoV-2019 Drug Development Team; Department of Biochemistry, Jomo Kenyatta University of Agriculture and Technology, Juja, Kenya; Research Scholar, Centre for Systems Biology and Bioinformatics Panjab University Chandigarh, INDIA; Research Associate, CSIR-IHBT, Palampur, H. P-176061; NIN-TATA Centre for excellence in public health nutrition, ICMR-National Institute of Nutrition, Jamai-Osmania (Post), Hyderabad, Telangana, INDIA; Bhaskaracharya College of Applied Sciences, University of Delhi, Delhi, INDIA

**Keywords:** SARS-CoV, SARS-COV-2, covid19, spike glycoproteins, phylogenetics, selection pressure, multi-epitope, vaccine, immune simulation

## Abstract

The recent outbreak of severe acute respiratory syndrome (SARS) coronavirus (CoV)-2 (SARS-CoV-2) causing coronavirus disease (covid19) has posed a great threat to human health. Previous outbreaks of SARS-CoV and Middle East respiratory Syndrome CoV (MERS-CoV) from the same CoV family had posed similar threat to human health and economic growth. To date, not even a single drug specific to any of these CoVs has been developed nor any anti-viral vaccine is available for the treatment of diseases caused by CoVs. Subunits present in spike glycoproteins of SARS-CoV and SARS-CoV-2 are involved in binding to human ACE2 Receptor which is the primary method of viral invasion. As it has been observed in the previous studies that there are very minor differences in the spike glycoproteins of SARS-CoV and SARS-CoV-2. SARS-CoV-2 has an additional furin cleavage site that makes it different from SARS-CoV (Walls et al., 2020). In this study, we have analyzed spike glycoproteins of SARS-CoV-2 and SARS-CoV phylogenetically and subjected them to selection pressure analysis. Selection pressure analysis has revealed some important sites in SARS-CoV-2 and SARS-CoV spike glycoproteins that might be involved in their pathogenicity. Further, we have developed a potential multi-epitope vaccine candidate against SARS-CoV-2 by analyzing its interactions with HLA-B*15:03 subtype. This vaccine consists of multiple T-helper (TH) cells, B-cells, and Cytotoxic T-cells (CTL) epitopes joined by linkers and an adjuvant to increase its immunogenicity. Conservation of selected epitopes in SARS, MERS, and human hosts, suggests that the designed vaccine could provide cross-protection. The vaccine is designed in silico by following a reverse vaccinology method acknowledging its antigenicity, immunogenicity, toxicity, and allergenicity. The vaccine candidate that we have designed as a result of this work shows promising result indicating its potential capability of simulating an immune response.

## Introduction

Coronavirus (CoV) is a pathogen that affects the human respiratory system. Previous severe outbreaks of CoVs have been emerged posing a great threat to public health. These outbreaks include Severe acute respiratory syndrome CoV (SARS-CoV) and Middle East Respiratory Syndrome CoV (MERS-CoV). Recently, in December 2019, a new type of coronavirus has emerged called SARS-CoV-2 or novel CoV-2019 (nCoV-2019) from Hubei province in China (Zhu et al., 2020). Since then, SARS-CoV-2 has posed a great threat to public health through coronavirus disease (covid19) (Pedersen & Ho, 2020; Velavan & Meyer, 2020). Due to the lack of proper antiviral treatment and vaccination, covid19 is spreading person to person at a very fast rate without being an air-borne disease (Jin et al., 2020; Rothan & Byrareddy, 2020).

There is no proper treatment available for covid19 except a few FDA approved drugs including Remdesivir (Agostini et al., 2018), Favipirar (“Favipiravir shows good clinical efficacy in treating COVID-19: official - Xinhua | English.news.cn,” 2020), Hydrochloroquine with Azithromycin (Gautret et al., 2020) have shown some excellent results on covid19 patients (Jean, Lee, & Hsueh, 2020). However, there is no such drug designed for the treatment of covid19 yet. Continuing outbreaks of CoVs pose a great threat to human health and still, there are no licensed vaccines for gaining protection against these CoVs are available yet. However, there have been some attempts to develop a few vaccines for SARS-CoV-2 including monoclonal antibody therapy (Shanmugaraj, Siriwattananon, Wangkanont, & Phoolcharoen, 2020) and a possible mRNA-SARS-CoV-2 vaccine (F. Wang, Kream, & Stefano, 2020). That calls for a desperate need for an anti-viral treatment and vaccine for SARS-CoV-2.

In this work, we have studied spike glycoproteins of SARS-CoV-2 as they have a furin cleavage site which is conserved amongst the 144 SARS-CoV-2 isolates (Zhou et al., 2020). This cleavage site located at the boundary of S1/S2 subunits differentiates SARS-CoV-2 from SARS-CoV and other related CoVs (Walls et al., 2020). Therefore, we have analyzed spike glycoproteins phylogenetically, subjected them to selection pressure analysis, and predicted epitopes for vaccine development. We have designed an in silico novel multi-epitope potential vaccine for SARS-CoV-2. The potential epitopes were predicted from spike glycoprotein sequences. These epitopes showing immunogenicity and antigenicity were selected. They were further filtered using molecular docking with HLA-B*15:03 allele as it has been reported to be responsive in inducing an immunogenic response effectively against SARS-CoV-2 (A. Nguyen et al., 2020). The interaction of constructed multi-epitope vaccine with HLA-B*15:03 allele was analyzed using molecular docking and showed various high-affinity bonds. Furthermore, the immune response of the candidate vaccine was simulated in silico and showed a significant immune response.

## Results

### Phylogenetic analysis

The constructed ML tree is shown in Figure 1. The phylogenetic tree of spike glycoproteins of SARS-CoV and SARS-CoV-2 showed distinct clades of these protein groups. However, the receptor-binding domain (RBD) in spike glycoprotein sequences (gi_1824676380_pdb_6W41_C and gi_1822249606_pdb_6M0J_E) of SARS-CoV-2 show larger branch lengths than the other sequences of the same. The low scoring sequences were eliminated from the rest of the sequences based on the phylogenetic tree. This resulted in a set of relevant spike glycoprotein sequences which were further subjected to selection analysis followed by antigenic sequence identification and in silico vaccine design.

**Figure 1.**
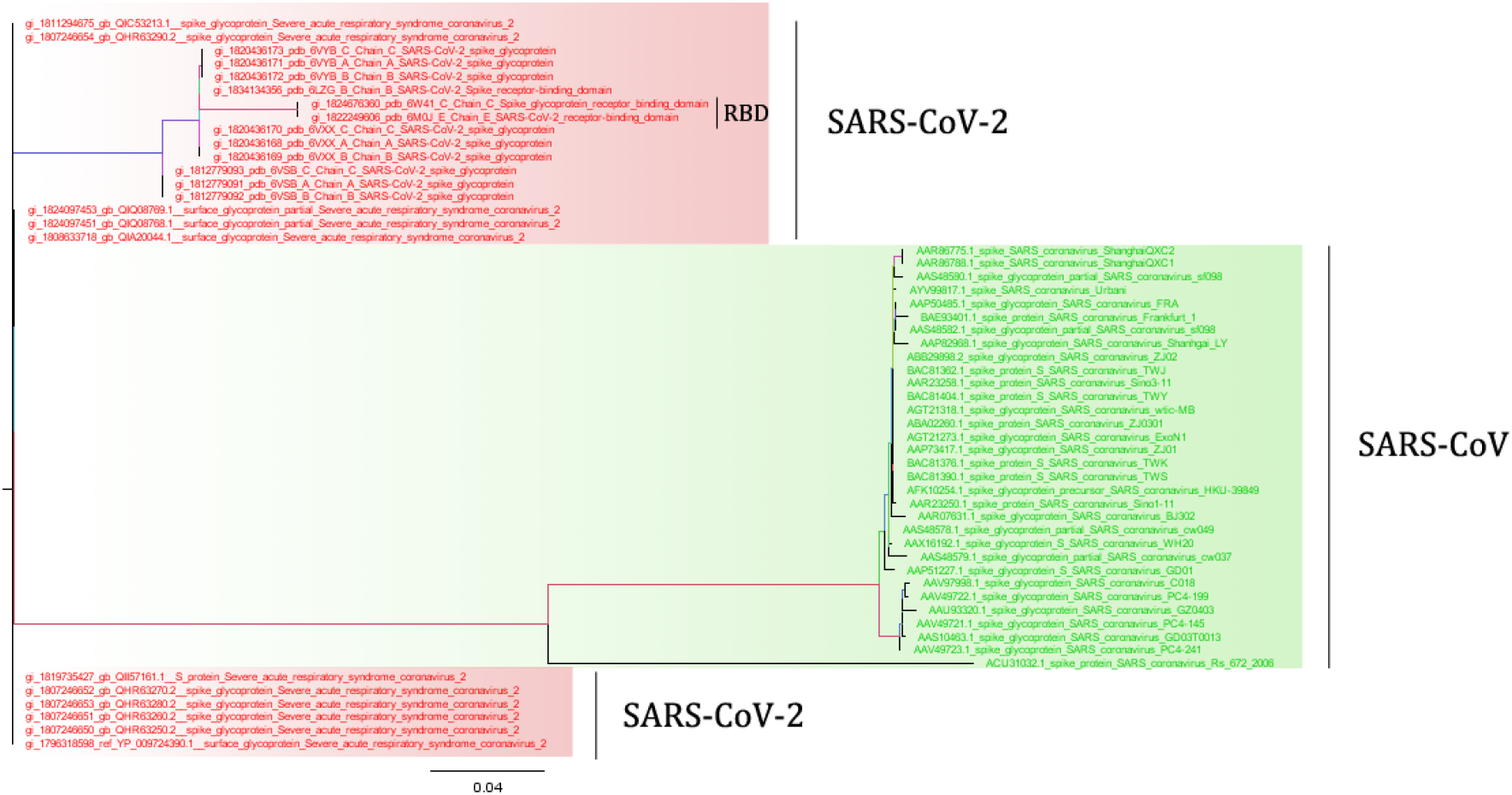
ML tree of spike glycoprotein sequences of SARS-CoV and SARS-CoV-2. *Red* sequences represent SARS-CoV-2 and *green* colored sequences show SARS-CoV spike glycoproteins.

### Selection pressure analysis

#### Spike glycoproteins of SARS-CoV-2 show positive selection

The selection pressure analysis of spike glycoprotein sequences of SARS-CoV-2 showed positive selection on a few sites. However, no single site has been identified which has experienced negative selection. Besides, BUSTED found no evidence of gene-wide episodic diversifying/positive selection. Three positively selected sites were identified by MEME at a p-value threshold of 0.05 and four such sites were identified by FUBAR (two sites coinciding with MEME) with a posterior probability of 0.9 (Additional file 1). Out of these four unique identified sites, two sites were found to be potentially relevant. These two sites, *Cys538* and *Thr549*, were mapped on the structure of SARS-CoV-2 spike glycoprotein (PDB ID: 6VXX) (Figure 2). As evident from the results shown in Figure 2, these sites were found to be present on β-sheets. The intermolecular interactions among β-sheets in a folded structure are considered important and have long been recognized (Nowick, 2008). They are involved in protein-protein interaction, protein quaternary structure, and protein aggregation. β-sheet interactions in some biological processes have been considered as potential targets for the treatment of diseases including cancer (J. Wang, Li, & Jasti, 2018) and AIDS (Li et al., 2013). Therefore, the two sites identified as positively selected sites may be relevant in spike glycoprotein structure stability and formation.

**Figure 2.**
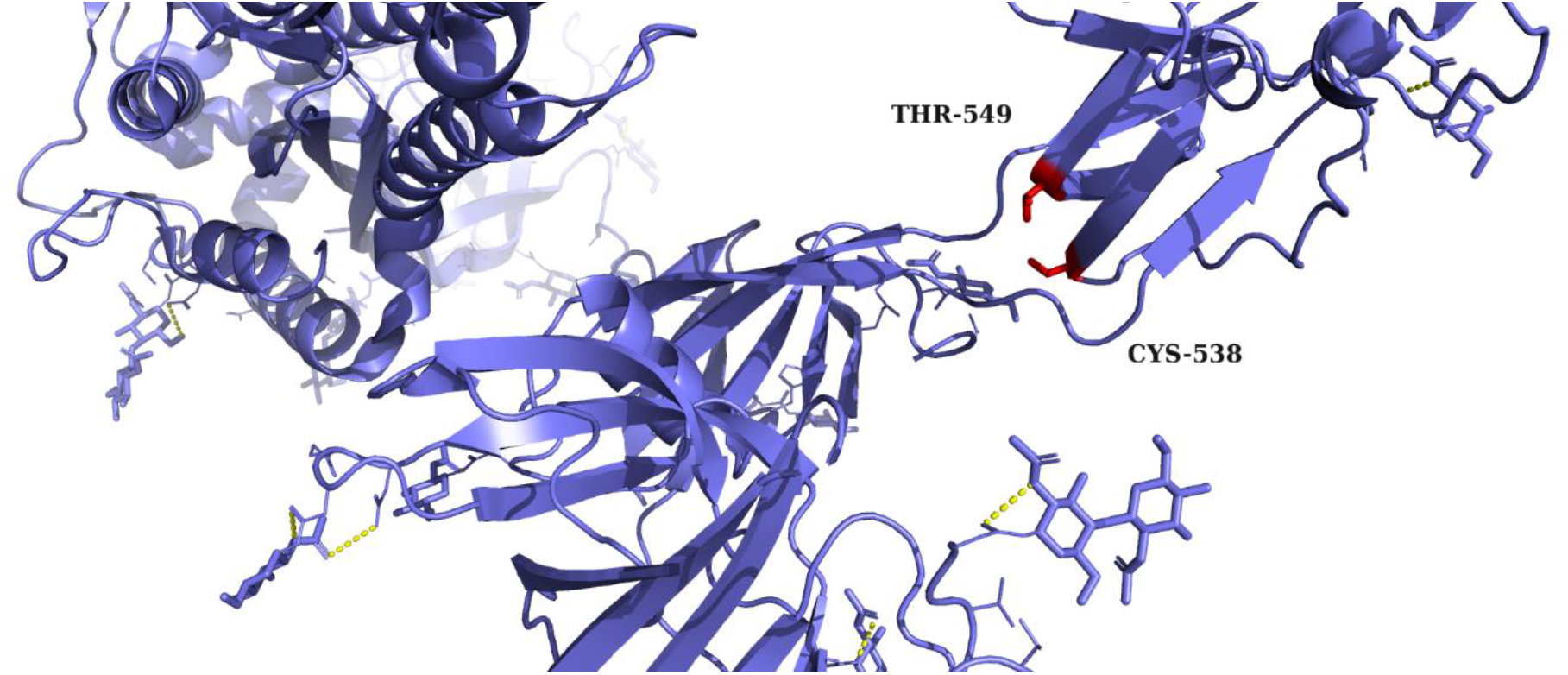
Two sites identified as positively selected mapped on crystal structure SARS-CoV-2 spike glycoprotein (PDB ID: 6VXX). The two identified sites are shown in *red* sticks.

#### Spike glycoproteins of SARS-CoV show positive selection

The selection pressure analysis of spike glycoprotein sequences of SARS-CoV showed positive selection on a larger number of sites as compared to that of SARS-CoV-2. However, like SARS-CoV-2 spike glycoproteins, no single site has been identified which has experienced negative selection. Unlike, SARS-CoV-2, BUSTED found evidence of gene-wide episodic diversifying selection at a p-value <=0.05 with synonymous rate variation. FUBAR has identified 2 sites of episodic positive selection with a posterior probability of 0.9 and MEME has identified 9 such sites at a p-value threshold of 0.05 (Additional file 2). These identified sites under positive selection were mapped on the crystal structure of spike glycoprotein of SARS-CoV (PDB ID: 5X58) to recognize their relevance. The well-studied structure of SARS-CoV spike glycoprotein (Walls et al., 2020) helped to locate the positively selected sites in different regions of the structure. The mapping revealed that a site *Thr244* lies in N-terminal domain (NTD), *Val594* and *Ala609* lie in Subdomain2 (SD2), *Thr743* and *Pro794* lie in Linker (L) and upstream helix (UH) region, *Leu803* lies in Fusion peptide (FP) region, and *Leu1031* lies in the central helix (CH), β-hairpin (BH), and Subdomain3 (SD3) region of SARS-CoV spike glycoprotein (Figure 3).

**Figure 3.**
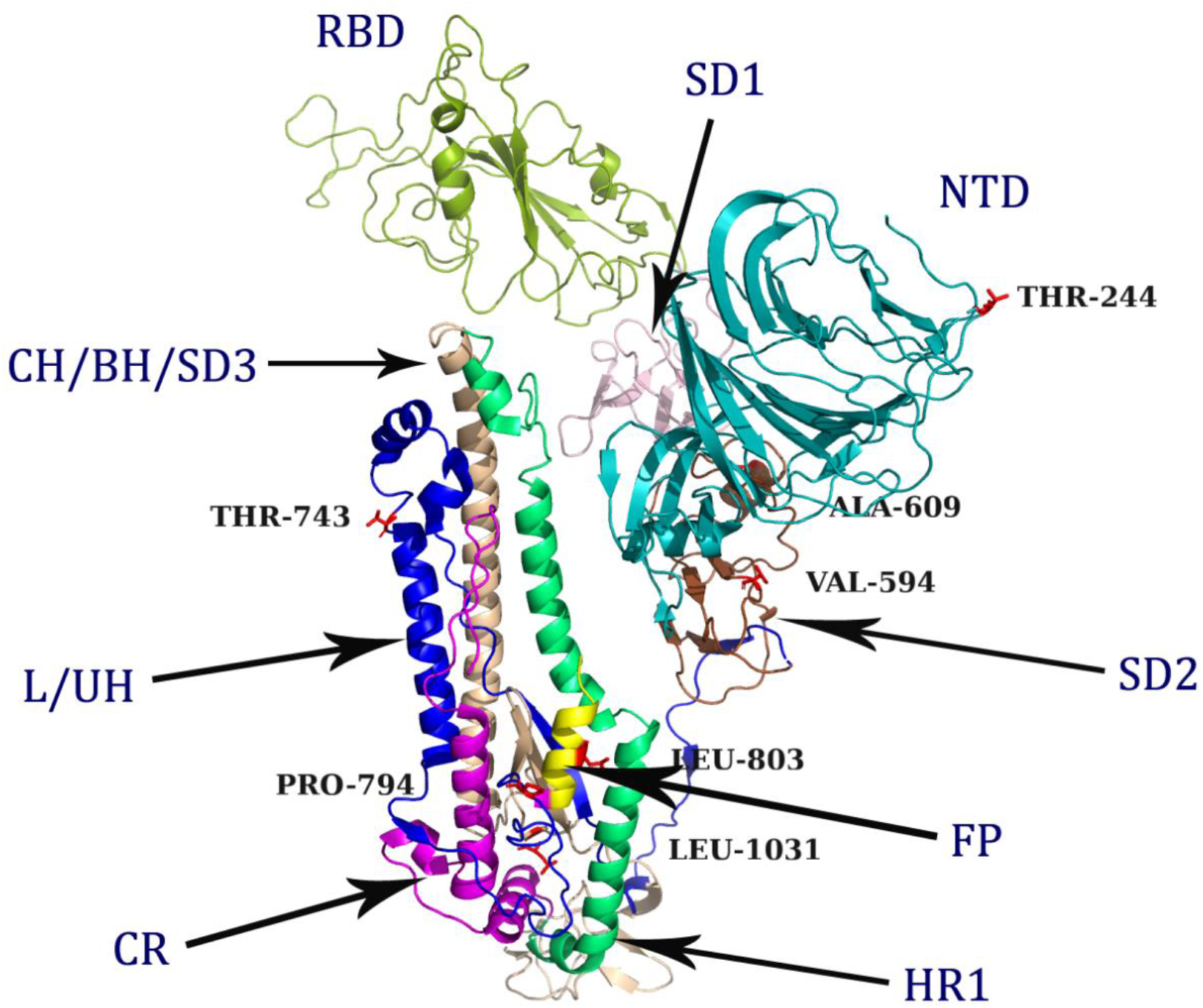
Identified positively selected sites (*red* sticks) mapped on crystal structure SARS-CoV spike glycoprotein (PDB ID: 5X58). Different colors on the structure depict different regions. NTD: N-terminal domain, RBD: Receptor binding domain, SD1: Subdomain1, SD2: Subdomain2, SD3: Subdomain3, CH: Central helix, BH: β-hairpin, L: Linker, UH: Upstream helix, FP: Fusion protein, HR1: Heptad repeat1.

### In silico design of multi-epitope vaccine

A complete scheme of in silico vaccine design is shown in Figure 4 and explained in the following sections.

**Figure 4.**
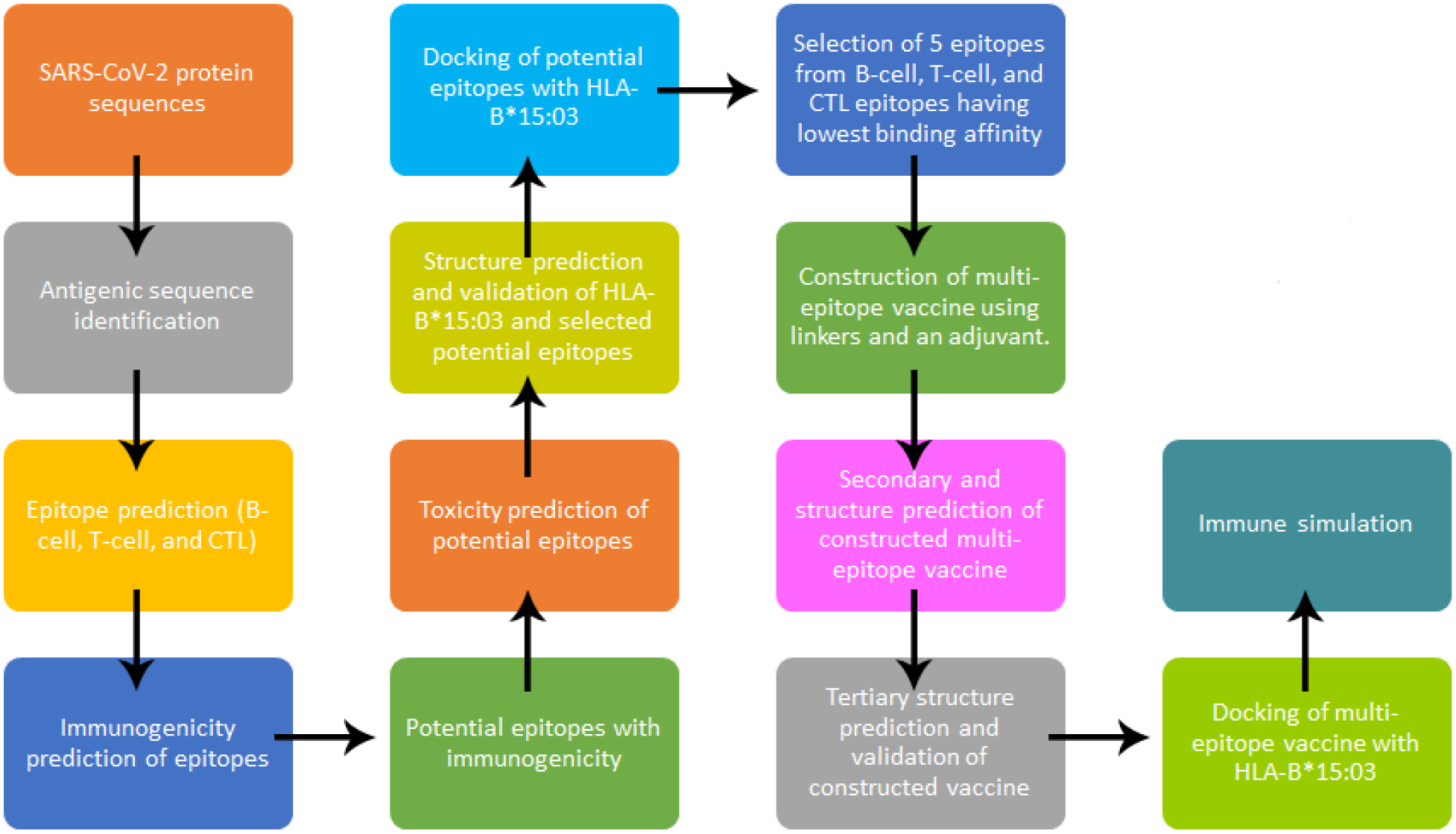
Schema of the methodology followed in the in silico design of a multi-epitope vaccine.

#### Antigenic sequences from SARS-CoV-2 spike glycoproteins

15 antigenic high scoring (threshold >0.5) sequences were obtained out of 23 sequences of SARS-CoV-2 spike glycoprotein submitted to the VaxiJen server. These selected sequences were used to identify T-cell, B-cell, and CTL epitopes.

#### Epitopes prediction from antigenic sequences of SARS-CoV-2 spike glycoproteins

The total number of T-cell, B-cell, and CTL predicted epitopes were 94, 89, and 47 respectively. The predicted epitopes were further analyzed for their immunogenicity. This provided 24, 15, and 7 T-cell, B-cell, and CTL epitopes with potential immunogenicity respectively. These epitope sequences were further subjected to toxicity analysis. All these resultant sequences showed non-toxic behavior and were subjected to further analyses.

#### Structure prediction of HLA-B*15:03 subtype

Swissmodel provides a quality estimation of predicted models using GMQE (Studer et al., 2020) and QMEAN score (Benkert, Biasini, & Schwede, 2011). GMQE (Global Model Quality Estimation) ranges between 0 and 1, a higher number implies higher reliability. QMEAN Z-score provides an estimate of experimental structures having a similar size. Positive values of QMEAN around zero indicate models with high quality. The HLA-B*15:03 subtype of HLA-B*15 allele with GMQE score of 0.78 and QMEAN of 0.81 (Figure 5A). Both the values indicate the predicted structure of the HLA-B*15:03 subtype is of high quality. The predicted structure was validated by plotting the Ramachandran plot. The plot showed 94.2% residues in most favored regions, 5% in additionally allowed regions, 0.8% in generously allowed regions, and 0% in disallowed regions (Figure 5B). This structure was used for molecular docking of the selected epitopes possessing potential immunogenicity.

**Figure 5.**
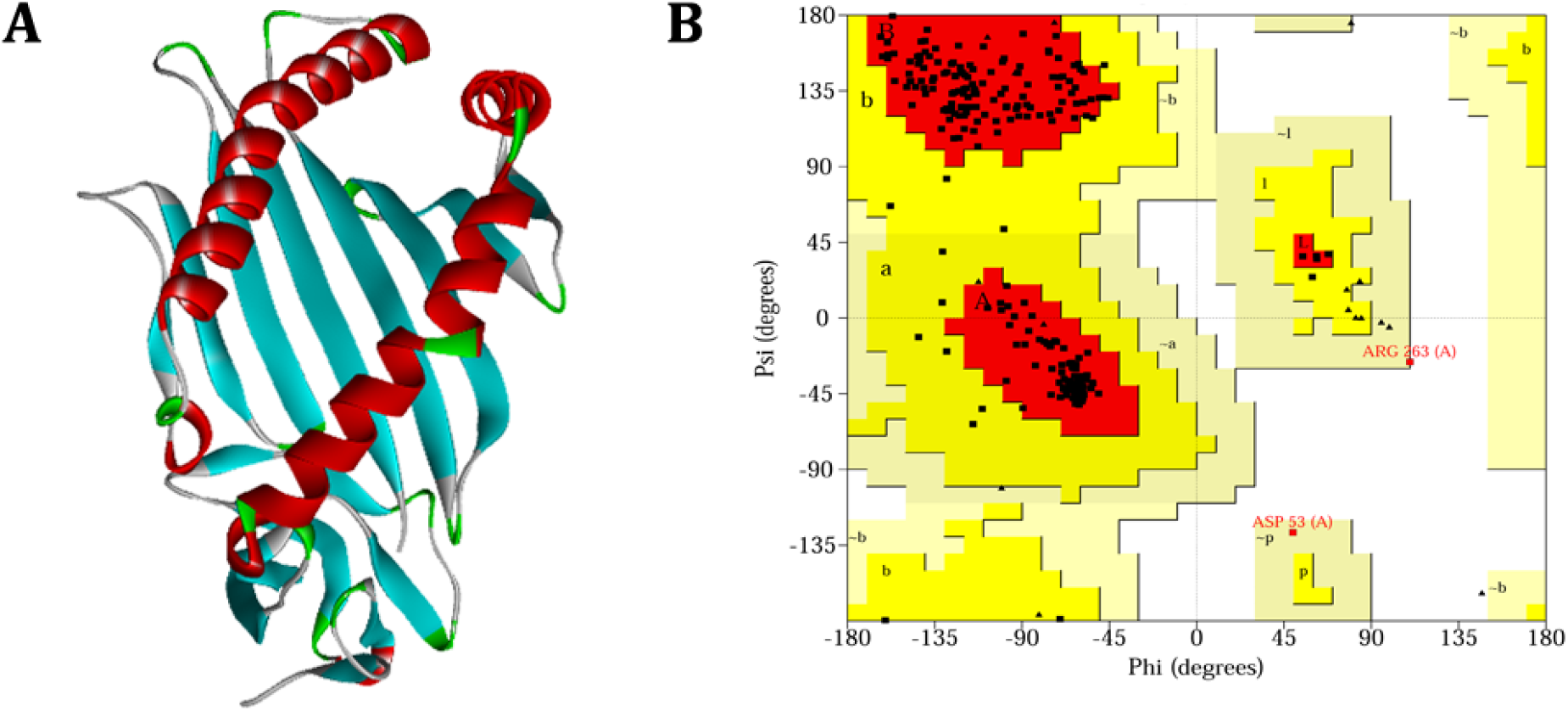
**A)** Predicted three-dimensional structure of HLA-B*15:03 allele. **B)** Ramachandran plot of the predicted structure of the HLA-B*15:03 allele.

#### Screening of potential epitopes for multi-epitope vaccine development

The selected epitope sequences were docked with HLA-B*15:03 subtype. The top five epitopes showing the lowest binding affinity were selected for further analyses. The binding affinities of the selected potential epitopes are shown in Table 1 along with their homology searched for MERS, SARS, and human host. According to the obtained docking results, these epitopes bound to the receptor effectively. This method was performed for each type of epitope (T-cell, B-cell, and CTL epitopes).

**Table 1.** Details of selected epitopes based on molecular docking with HLA-B*15:03.

| Type of epitope | Sequence ID | Sequence | Homology | No of amino acid residue | Binding affinity (kcal/mol) |
| --- | --- | --- | --- | --- | --- |
| T-Cell | QIK02137.1 ORF8_1 | IQYIDIGNY | 100 | 9 | -6.5 |
|  | YP_009742610.1 nsp3_7 | VQTIEVNSF | 100 | 9 | -6.5 |
|  | YP_009742610.1 nsp3_24 | LVAEWFLAY | 100 | 9 | -6.2 |
|  | QJD49277.1 ORF10_2 | YINVFAFPF | 55 | 9 | -6.1 |
|  | YP_009742610.1 nsp3_45 | VVVNAANVY | 100 | 9 | -6.1 |
| B-Cell | QIK02137.1_5 | EDFLEY | 100 | 6 | -8.8 |
|  | gi_1822249606_pdb_6M0J_E_8 | CVNFHH | 100 | 6 | -8.8 |
|  | gi_1831503521_pdb_6YLA_A_5 | SYGFQPTNGVGYQ | 92 | 13 | -8.1 |
|  | gi_1822249606_pdb_6M0J_E_1 | ATRFAS | 66 | 6 | -7.9 |
|  | YP_009742610.1_22 | KTVGELGD | 100 | 8 | -7.8 |
| CTL | gi_1820435683_3 | NLCPFGEVF | 100 | 9 | -7.1 |
|  | YP_009742610.1 nsp3_1 | FGDDTVIEV | 100 | 9 | -5.9 |
|  | YP_009742610.1 nsp3_3 | FSYFAVHFI | 100 | 9 | -5.1 |
|  | YP_009742610.1 nsp3_4 | APYIVGDVV | 100 | 9 | -6.2 |
|  | YP_009742616.1 nsp9_1 | KGGRFVLAL | 100 | 9 | -6.5 |

#### In silico design of multi-epitope vaccine

The designed multi-epitope vaccine consisted of 699 amino acid residues including linkers, adjuvant, and a 6x-His tag added at the C-terminus for purification purposes (Figure 6). This potential vaccine is non-allergen. Molecular weight is 75143.19, theoretical pI is 5.36, the Grand average of hydropathicity (GRAVY) is 0.016, the aliphatic index is 90.89, and instability index is 35.14 classifying potential vaccine as stable. Predicted secondary structure of the multi-epitope vaccine candidate consists of 34% alpha-helices, 28% beta-sheets, and 36% coils (Figure 7). The predicted solubility of the multi-epitope vaccine was a little below average value (0.45) (Figure 8). It implies that the multi-epitope is less soluble in *E. coli* protein.

**Figure 6.**
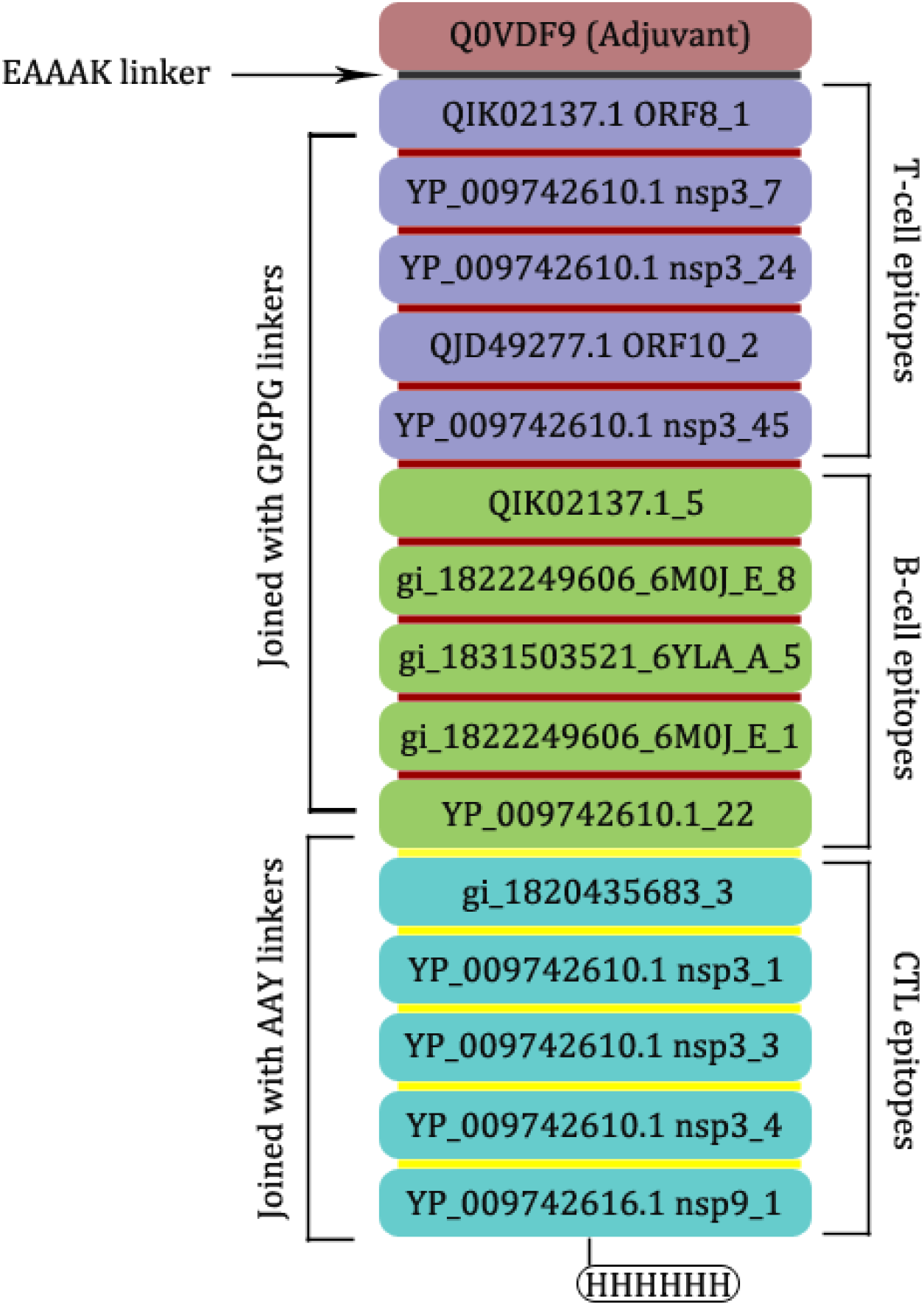
Schematic presentation of the designed 699 amino acids long multi-epitope vaccine. An adjuvant is added at the amino-terminal with the help of the EAAAK linker (grey). T_h_ cell epitopes and B-cell epitopes were joined using GPGPG linkers (red) and CTL epitopes were joined using AAY linkers (Yellow). TH cell, B-cell, and CTL epitopes are depicted with *light blue, light green*, and *sky blue* colors respectively.

**Figure 7.**
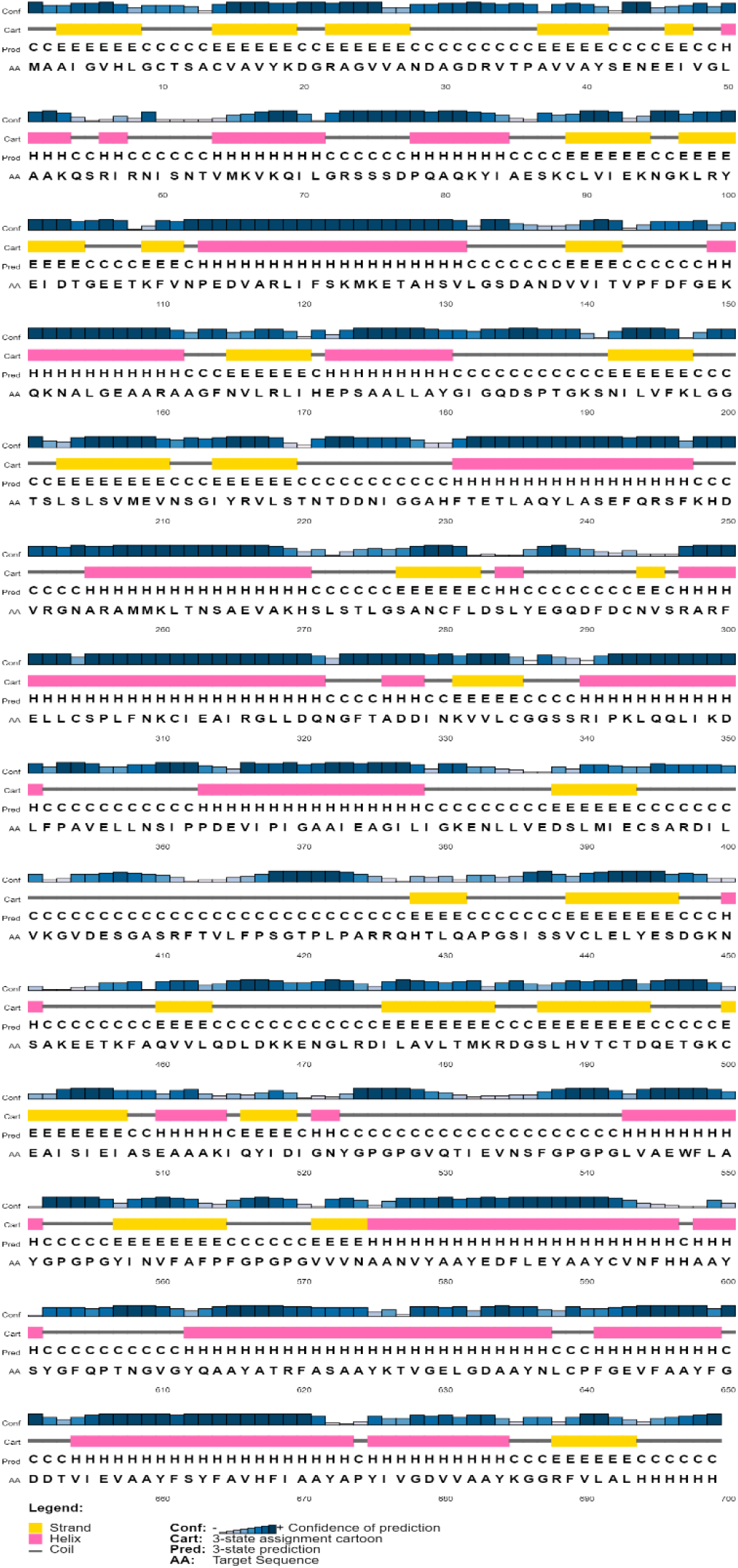
Predicted secondary structure of the multi-epitope vaccine.

**Figure 8.**
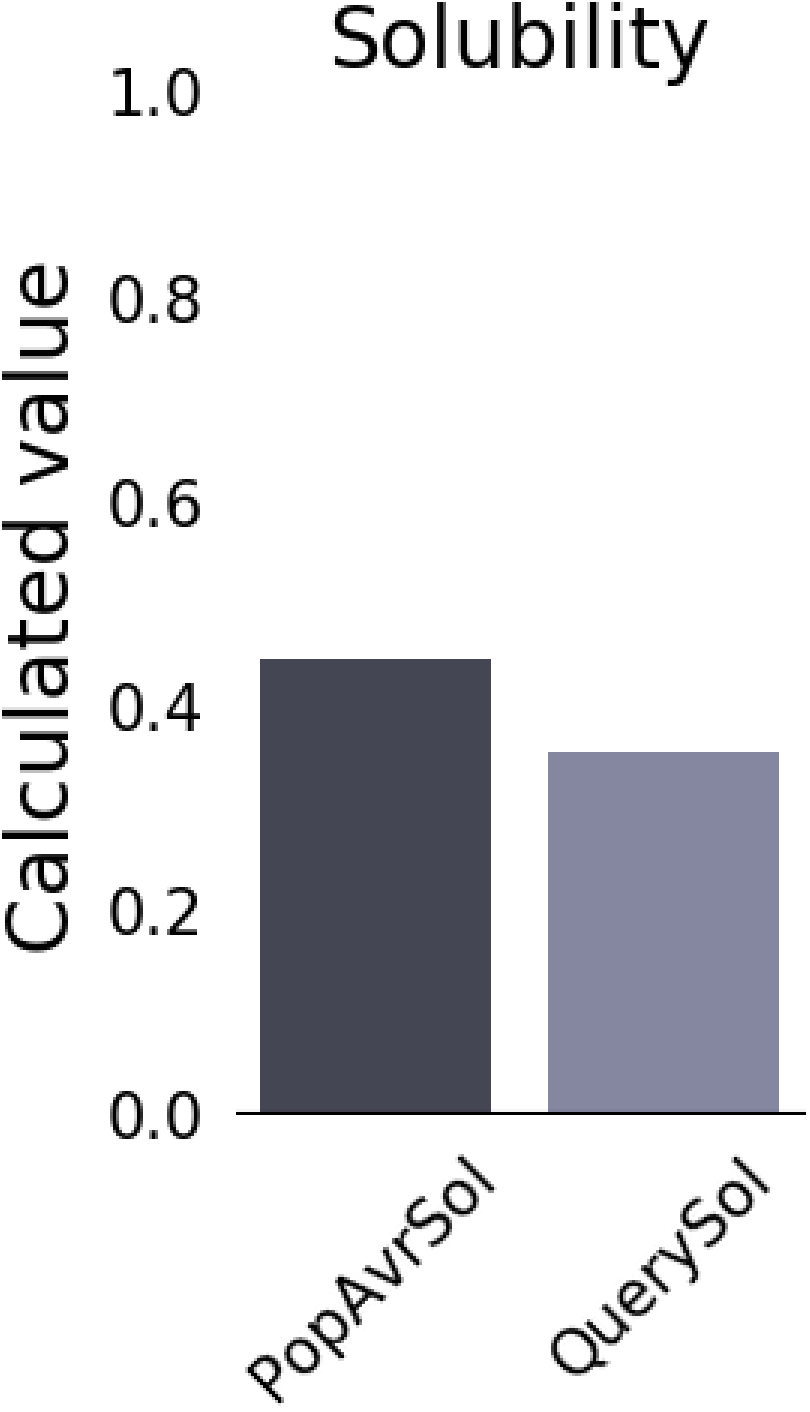
Predicted solubility of the designed multi-epitope vaccine. QuerySol represents the vaccine and PopAvrSol represents the average solubility (0.45) of *E. coli* protein.

I-Tasser predicted five models, out of which model 1 was selected having a C-score of - 2.03 (Figure 9A). The predicted structure was validated by plotting the Ramachandran plot.

**Figure 9.**
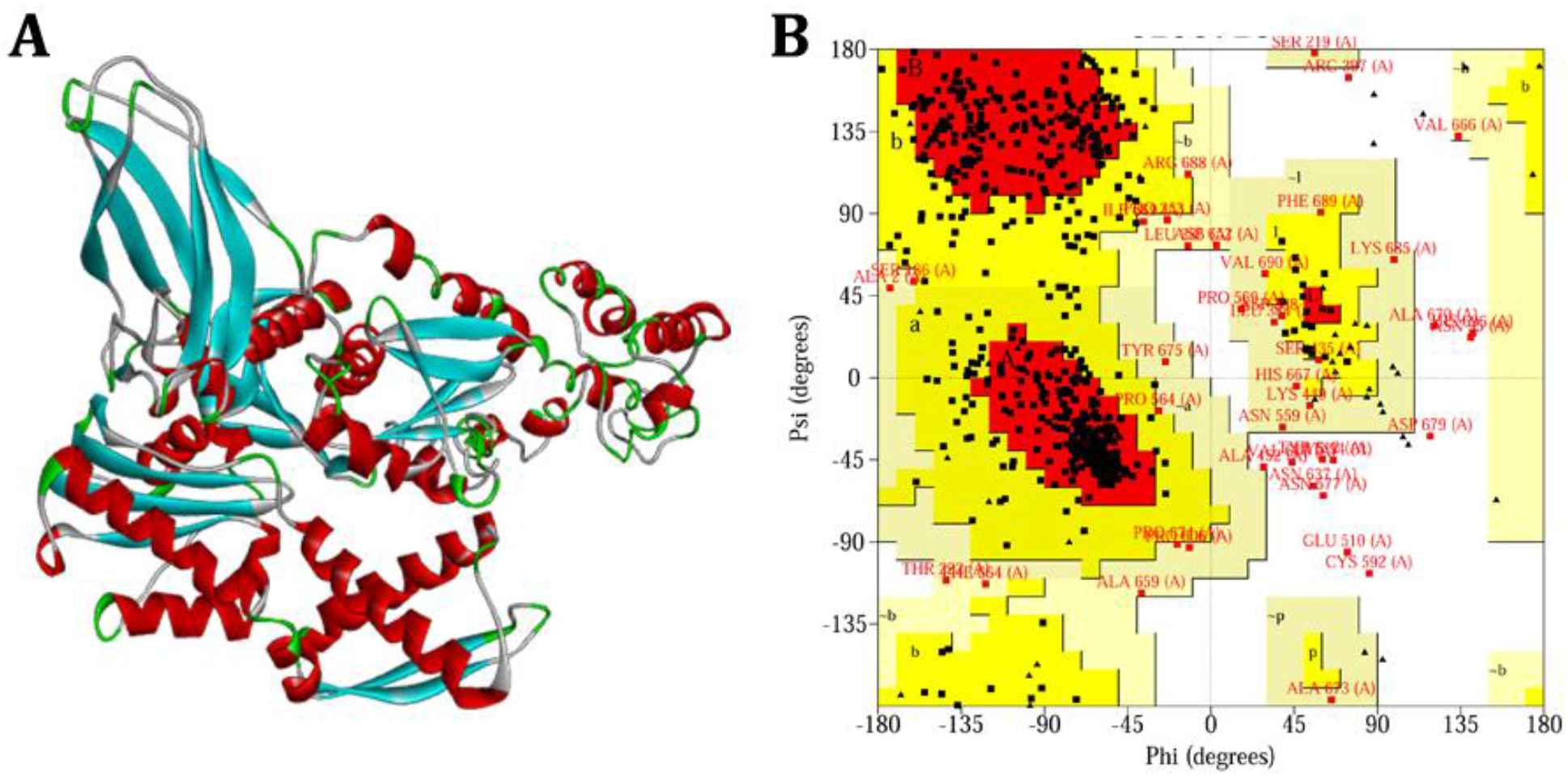
**A)** Predicted three-dimensional structure of the multi-epitope vaccine. **B)** Ramachandran plot of the predicted structure of the multi-epitope vaccine.

The plot showed 69% residues in most favored regions, 25.2% residues in additionally allowed regions, 3.6% residues in generously allowed regions, and 2.1% residues in disallowed regions (Figure 9B).

#### multi-epitope vaccine binds effectively with HLA-B*15:03

The top-ranking pose was selected from HADDOCK showing Z-score of −1.9, the most negative value amongst all poses (Z-score more negative is better). Docked complex of multi-epitope vaccine candidate and HLA-B*15:03 subtype was analyzed using the PRODIGY server and later by Pymol. The binding affinity was found to be −13.5 kcal/mol and dissociation constant (Kd) was 1.2E-10 M at 25°C. A Pymol script generated by PRODIGY helped to visualize the interacting residues between the multi-epitope vaccine candidate and the predicted HLA-B*15:03 subtype structure (Figure 10A). The number of contacts made at the interface (ICs) per property include charged-charged: 3, charged-polar: 3, charged-apolar: 41, polar-polar: 5, polar-apolar: 25, and apolar-apolar: 24. A list of all interacting residues is provided in Table 2. Amongst them, the closely interacting residues include *Ile90, Lys92, Glu100, Ser155, Ala174, Gln179, Arg181*, and *Arg194* (Figure 10B). The molecular docking of candidate multi-epitope vaccine with HLA-B*15:03 suggests that this designed vaccine may be a good candidate against covid19.

**Table 2.** All interacting residues of HLA-B*15:03 and multi-epitope vaccine obtained after molecular docking.

| HLA-B*15:03 |  | Multi-epitope vaccine |  |
| --- | --- | --- | --- |
| Interacting residues | #Amino acid | Interacting residues | #Amino acid |
| LYS | 92 | TYR | 557 |
| GLN | 89 | PHE | 565 |
| LYS | 170 | TYR | 600 |
| ALA | 174 | TYR | 600 |
| ILE | 90 | PRO | 567 |
| ARG | 175 | ILE | 669 |
| ASP | 85 | PRO | 569 |
| SER | 101 | ASN | 559 |
| GLN | 179 | VAL | 560 |
| GLU | 178 | ALA | 665 |
| THR | 167 | TYR | 602 |
| SER | 155 | PHE | 664 |
| THR | 93 | PHE | 561 |
| ARG | 86 | PHE | 645 |
| ARG | 175 | PHE | 664 |
| GLU | 190 | ASP | 652 |
| GLU | 82 | PHE | 645 |
| GLN | 96 | TYR | 557 |
| ILE | 166 | TYR | 602 |
| GLN | 179 | ALA | 562 |
| TRP | 191 | THR | 653 |
| TYR | 83 | PHE | 645 |
| ASP | 85 | PRO | 567 |
| TYR | 183 | THR | 653 |
| GLY | 80 | TYR | 648 |
| ARG | 86 | PHE | 641 |
| ASN | 104 | ASN | 559 |
| ARG | 175 | PHE | 661 |
| GLU | 82 | PRO | 640 |
| GLN | 89 | GLY | 568 |
| ALA | 173 | TYR | 602 |
| GLU | 87 | PHE | 645 |
| THR | 93 | ASN | 559 |
| THR | 97 | ASN | 559 |
| THR | 97 | TYR | 557 |
| ALA | 177 | PHE | 664 |
| GLU | 185 | TYR | 660 |
| GLN | 89 | PHE | 561 |
| GLU | 178 | TYR | 660 |
| GLU | 176 | VAL | 560 |
| ARG | 169 | TYR | 602 |
| ASN | 94 | TYR | 557 |
| GLU | 82 | PHE | 641 |
| ALA | 182 | ILE | 655 |
| ALA | 177 | TYR | 660 |
| ALA | 174 | VAL | 560 |
| ARG | 86 | PRO | 567 |
| GLU | 82 | PRO | 567 |
| GLN | 89 | PRO | 567 |
| GLU | 100 | ASN | 559 |
| ARG | 194 | ASP | 651 |
| ARG | 181 | TYR | 663 |
| GLY | 80 | PHE | 649 |
| TYR | 83 | PRO | 567 |
| GLU | 178 | VAL | 657 |
| TRP | 191 | ASP | 651 |
| LEU | 187 | ASP | 652 |
| GLN | 179 | PHE | 561 |
| ARG | 181 | PHE | 664 |
| ASP | 85 | GLY | 568 |
| ALA | 173 | PHE | 604 |
| GLU | 178 | PHE | 664 |
| ARG | 175 | ALA | 670 |
| TYR | 183 | TYR | 660 |
| ASN | 94 | ASN | 559 |
| TRP | 191 | PHE | 645 |
| GLU | 190 | ASP | 651 |
| LEU | 180 | VAL | 560 |
| LYS | 170 | TYR | 602 |
| ARG | 194 | PHE | 649 |
| PRO | 81 | TYR | 648 |
| THR | 97 | VAL | 560 |
| ARG | 86 | GLY | 568 |
| GLU | 79 | PHE | 649 |
| GLN | 89 | TYR | 557 |
| GLN | 89 | GLY | 566 |
| THR | 93 | TYR | 557 |
| ARG | 181 | TYR | 660 |
| LEU | 187 | THR | 653 |
| ARG | 175 | ALA | 682 |
| GLN | 179 | VAL | 657 |
| ARG | 194 | GLY | 650 |
| GLU | 82 | VAL | 644 |
| LYS | 170 | ASN | 559 |
| GLN | 89 | PRO | 569 |
| GLU | 100 | GLY | 556 |
| ARG | 175 | ALA | 665 |
| GLU | 178 | ILE | 669 |
| PRO | 81 | PHE | 649 |
| ALA | 182 | TYR | 660 |
| ARG | 86 | THR | 653 |
| LEU | 187 | ILE | 655 |
| TRP | 191 | PHE | 649 |
| GLN | 179 | ILE | 655 |
| TYR | 83 | PHE | 649 |
| TYR | 183 | ILE | 655 |
| GLU | 82 | PHE | 649 |
| LEU | 187 | ASP | 651 |
| GLN | 165 | TYR | 602 |
| GLN | 179 | LEU | 549 |
| ALA | 174 | LEU | 549 |

**Figure 10.**
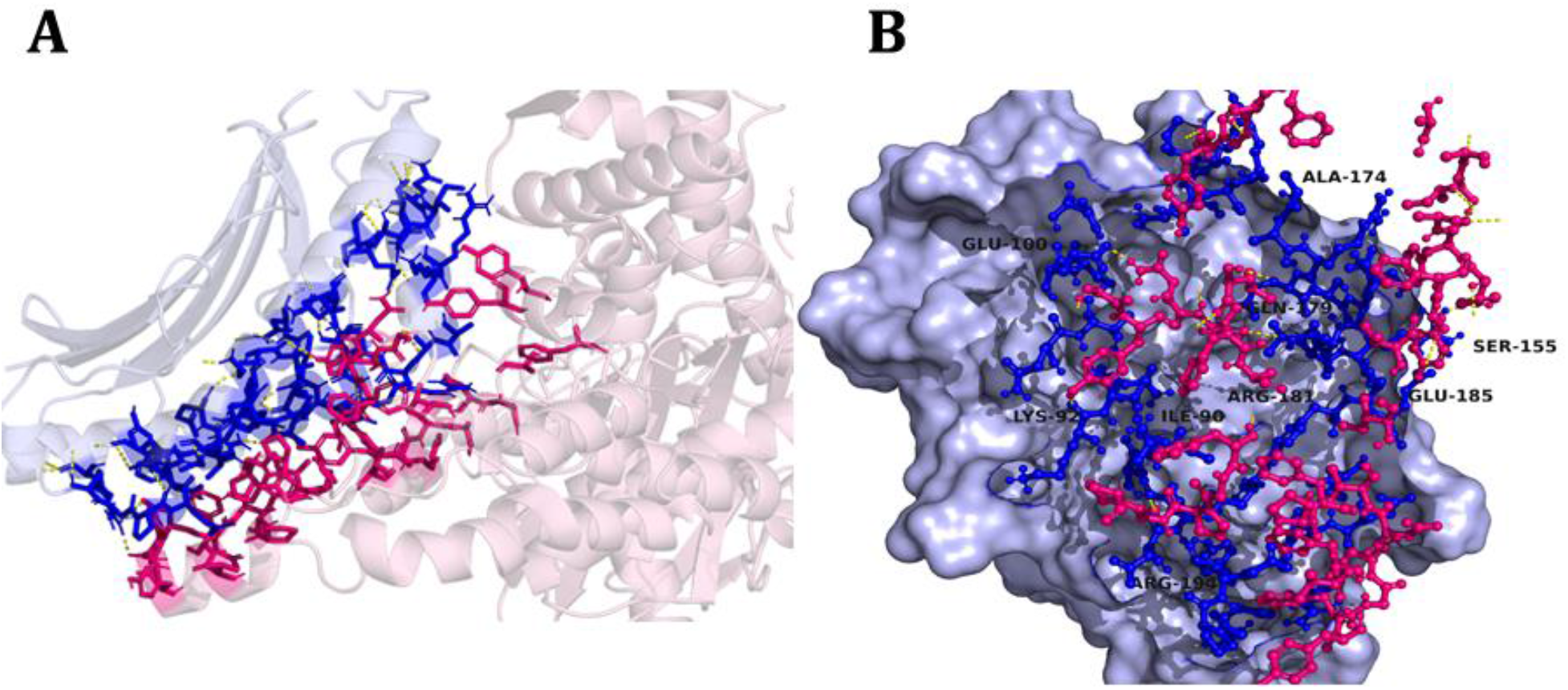
Interaction of designed multi-epitope vaccine with HLA-B*15:03 allele. **A)** interfaces of multi-epitope vaccine (*magenta*) and HLA-B*15:03 (*blue*) interacting with each other generated using Pymol script. **B)** Most prominent interacting residues of HLA-B*15:03 shown in *blue* color-forming bonds with the residues of multi-epitope vaccine (*magenta*).

#### In silico Immune simulation of multi-epitope vaccine shows a significant immune response

##### T helper cells (CD4)

After day 0, total TH cells (CD4) were noticed to increase from 1300 to reach a maximum of 6000 then rise again after the second injection at day21 to reach 10000 on day 27. In the meantime, TH memory cells level increased until it reached 4000 on day 21 then jumped to reach 9000 cells and fell gradually (Figure 11A). Most of these cells were in an active state reaching a level of 3000 on day21 then increased again to 5000 cells on day 30 after vaccination while few of them became tolerant anergic in the first week (Figure 11B). Active regulatory TH cells started to rise from 20 at the beginning of vaccination to a level of 160 after 2 days then fell gradually (Figure 11C). TH1 (shown in plot as Th1) level increased to reach a level of 50000 at day 21 then kept rising to reach 90000 cells by the day 27 (Figure 11D) showing 100% till the end of the third month (Figure 11E) while the T regulatory (TR) cells did not exceed the number of 160 cells.

**Figure 11.**
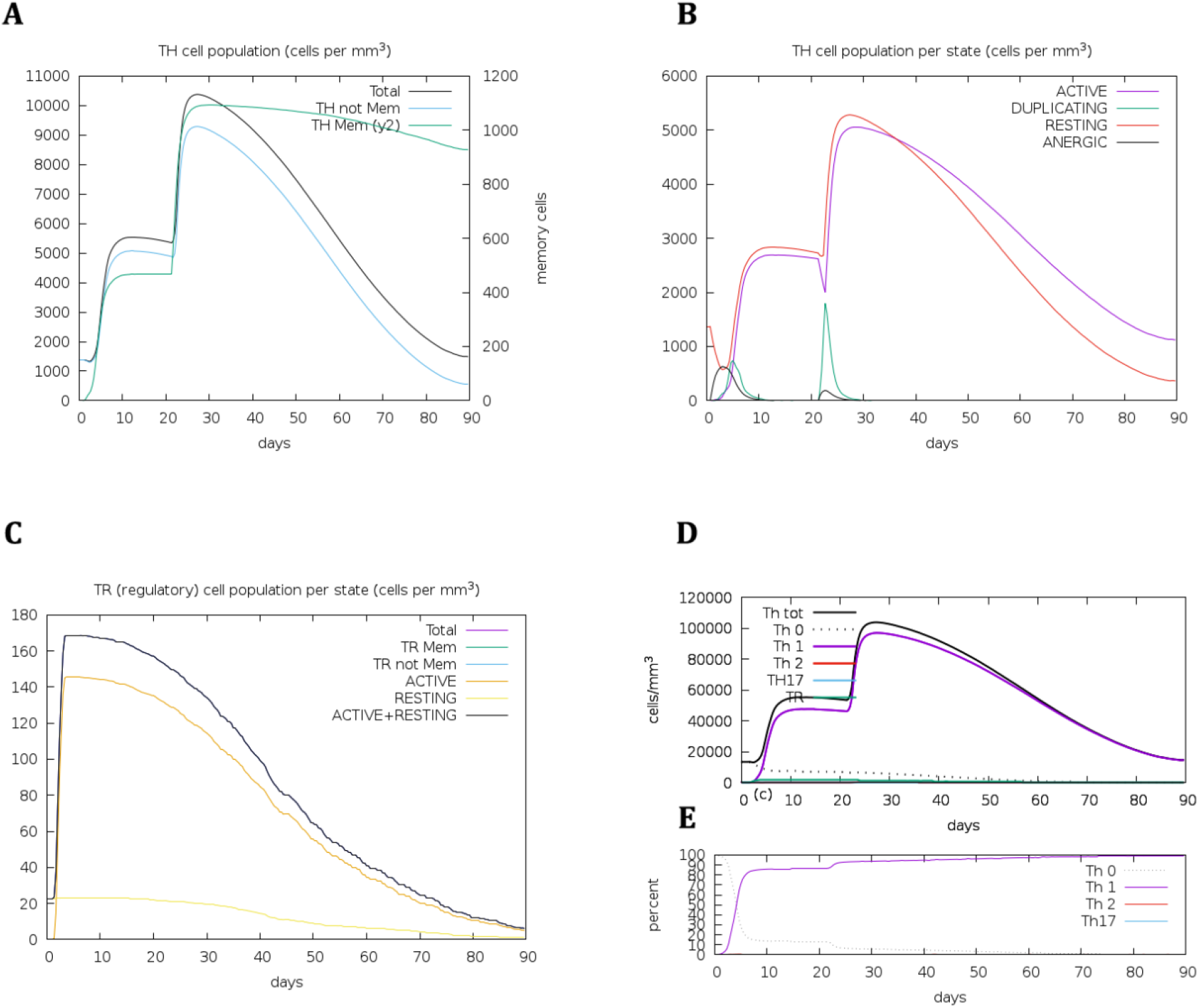
Distribution of TH cells during vaccination. **A)** Distribution of memory and non-memory cells. **B)** Four state distribution of TH cells. **C)** CD4 T-regulatory lymphocytes count with both total memory and per entity-state counts. **D)** Different stages of TH cells. **E)** Percentage of TH cells distribution for three months. TH: T-helper cell, Mem: memory cell, TR: T-helper regulatory cells.

##### CTL (CD8)

Non-memory CTL (CD8) can be seen increasing from below 1100 after day 0 to reach a maximum level of 1130 number of cells on day 23, then falling again only to rise at day 27 (6 days after the second injection) (Figure 12 A). Meanwhile, CTL memory cells continued to rise until they reached to a number of 1130 cells. CTL active cells were predicted to increase after day 0 to reach a level of 900 at day 40 before it started to fall. Tolerant anergic cells did not exceed the level of 600 throughout the simulation (Figure 12B). Dendritic cells (DC) can either present antigen to MHC1 or MHC2. Here, around 200 DCs are presenting antigens to MHC2 in comparison to only 10 DC cells presenting to MHC1 (Figure 12C).

**Figure 12.**
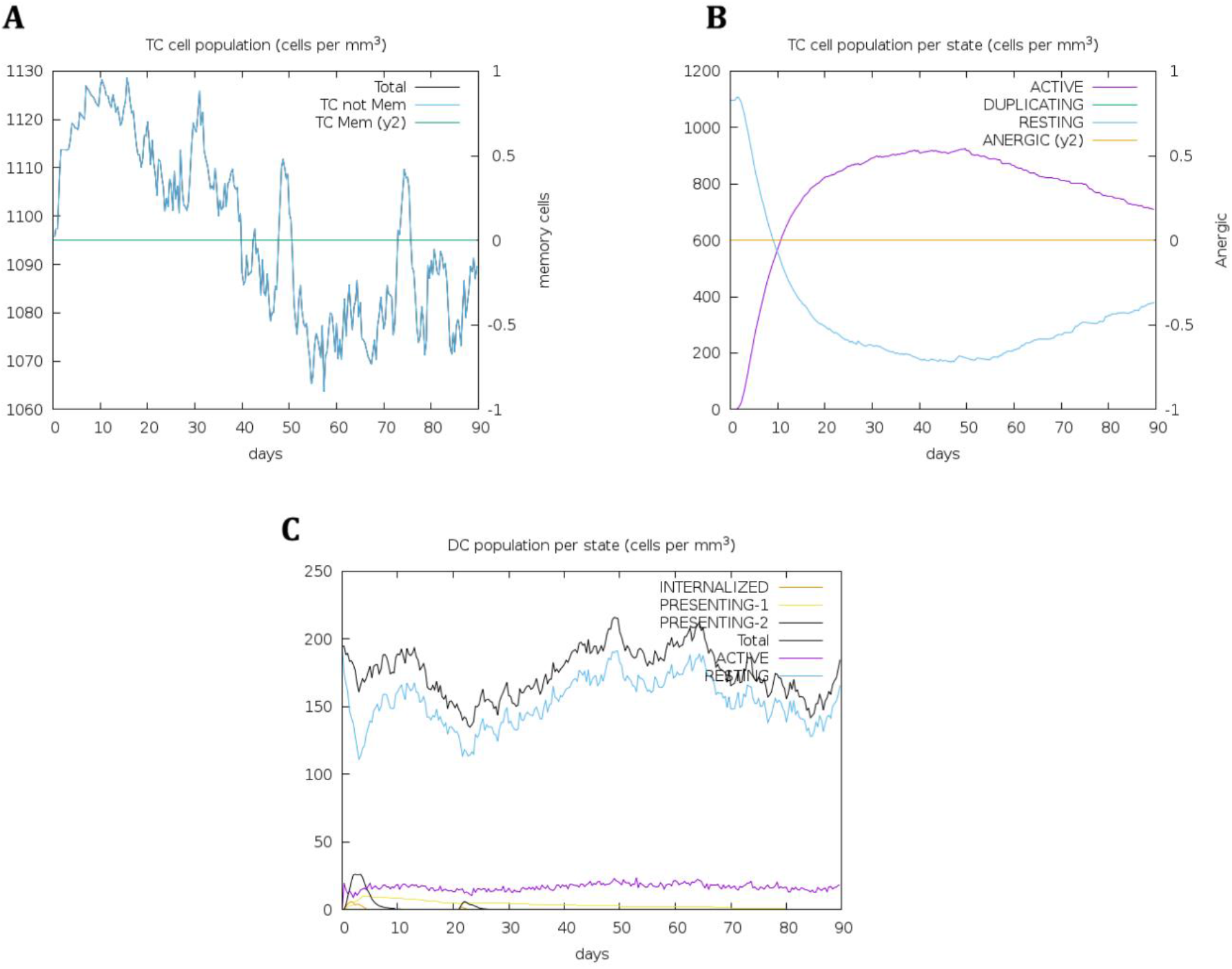
Distribution of CTL during vaccination. **A)** Distribution of memory and non-memory cells. **B)** Four state distribution of TC cells. **C)** Different states of dendritic cells (DC) including resting, active, and presenting states. The resting state represents the cells not presented to the antigen and anergic state refers to tolerance of T-cells to the antigen due to repeated exposures. TC: T-cytotoxic cell, Mem: Memory cells, and DC: dendritic cells.

##### B-cells and antibodies

The total B-cell number increased from 0 on day 1 to near 700 on day 21 before slowly falling again (Figure 13A). Moreover, antibodies production was noticed to occur almost five days after the beginning of infection with an earlier rise in Immunoglobulin-M (IgM) followed by the rise of Immunoglobulin-G (IgG). Both were noticed to fall until day 20, before rising again after the second injection at day 21 (Figure 13B).

**Figure 13.**
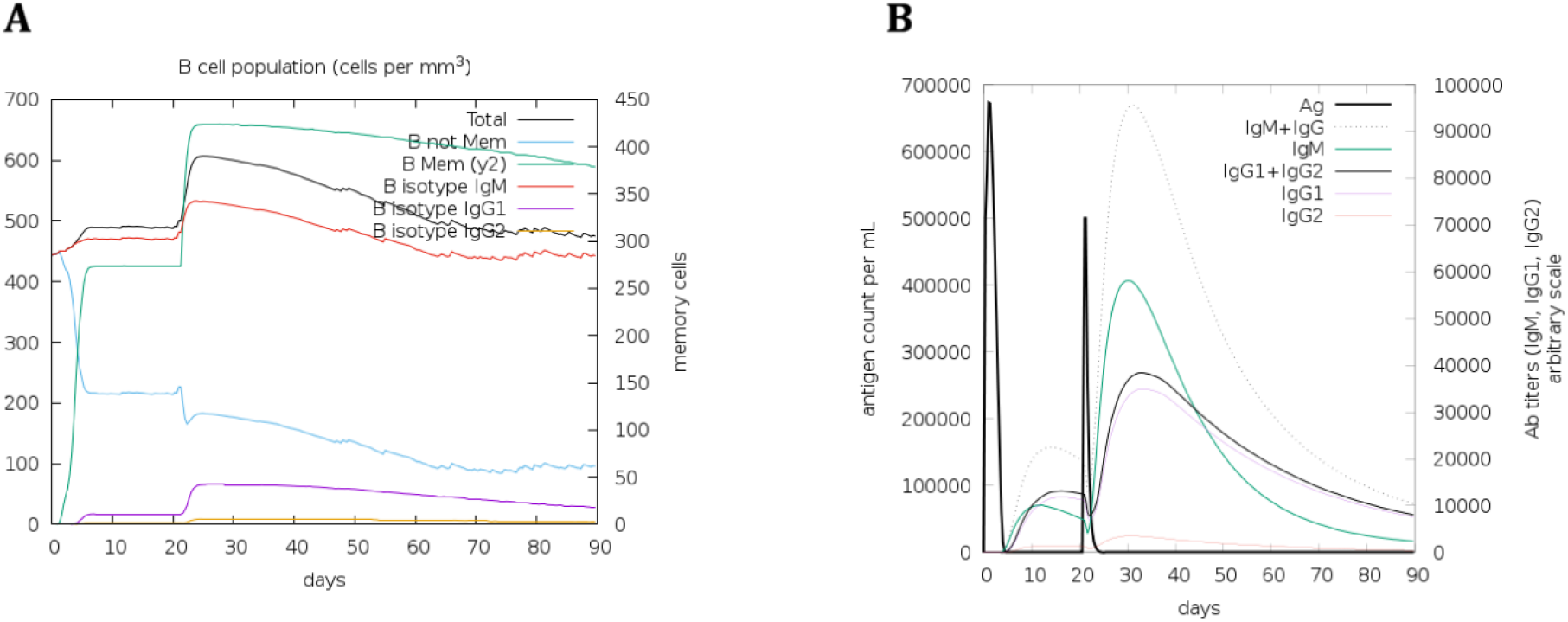
Secondary antibodies response simulation during vaccination. **A)** formation of different types of B-cells. **B)** earlier increase in IgM as compared to IgG.

##### Cytokines, Interleukins, and Natural killer (NK) cells

Cytokines are necessary for successful vaccination. Here, for our designed vaccine candidate, all cytokines started to increase after day 0 to reach a maximum level at day 5 and 7, then started to fall, only to rise again after the second vaccine injection at day 21 to reach similar levels except for Interferon-ɣ (IFN-ɣ) whose subsequent rise is only up to 1250 in comparison to 420000 in the first injection reaction. Special focus was on the response of Interleukin-2 (IL-2), where a higher level (450000) of IL-2 was noticed after the second injection (Figure 14A).

**Figure 14.**
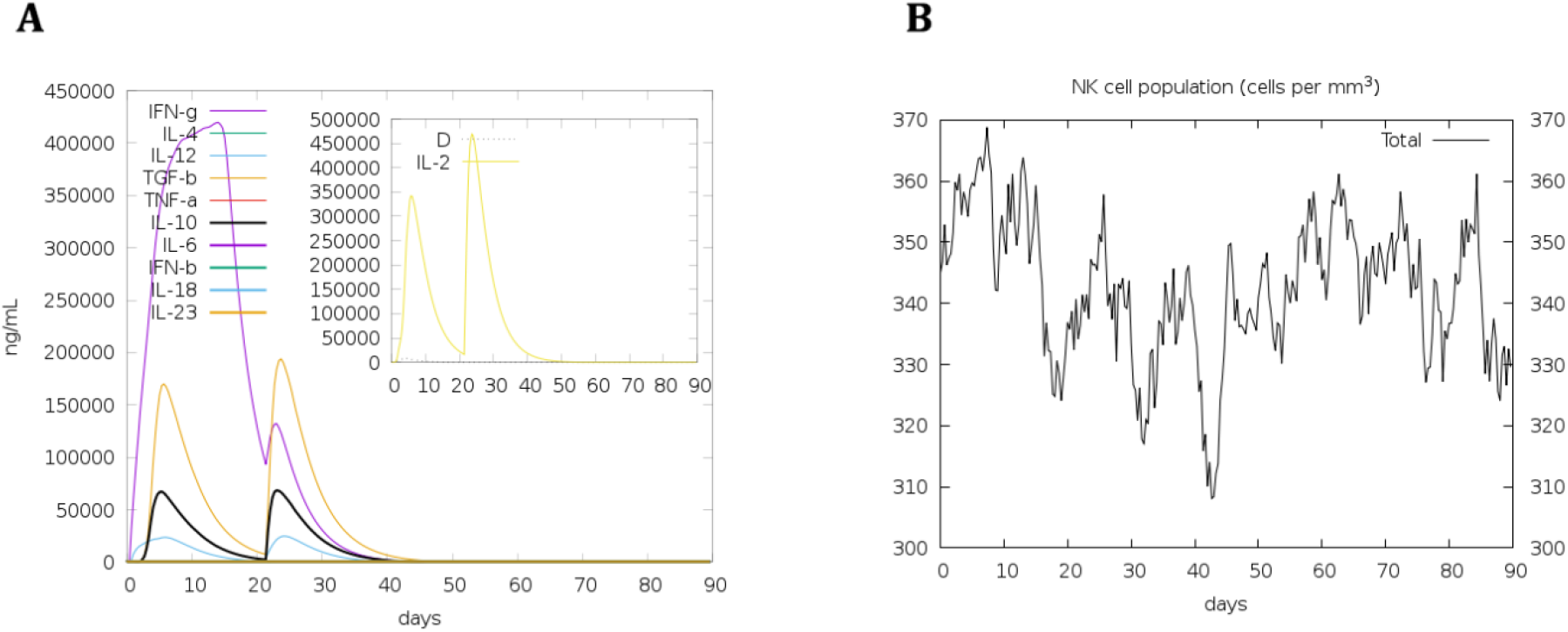
Cytokines, IL-2, and NK cells simulated reaction to vaccination. **A)** different levels of cytokines through time with a focus on interleukin 2 (IL-2). **B)** Distributions of NK-cells after vaccination showing high fluctuation levels between 310 and 375 cells.

NK cells are an integral part of the early response to the virus. The in silico simulation of vaccine candidate showed high fluctuations in the population of NK cells throughout the 90 days beginning from a level of 345 cells then fluctuated to reach a level around 330 by the end of third month (Figure 14B).

## Conclusion

In this study, we have studied spike glycoproteins of SARS-CoV and SARS-CoV-2 phylogenetically and designed a multi-epitope vaccine for covid19. The selection pressure analysis on spike glycoproteins revealed several important sites that have undergone positive selection. Interestingly, no negative selection was found in both types of CoVs. As negative selection selectively removes the harmful variants, it indicates that the selection process in SARS-CoVs has not been stabilized yet. Additionally, it also suggests that the negative selection in SARS-CoVs is quite weak that is allowing the widespread of deleterious mutations in the genome. This can eventually lead to the back mutations which may potentially act as one of the contributing factor in maintaining the genomic integrity of the virus (Loewe, 2008). Besides, unlike SARS-CoV, the gene-wide episodic diversifying selection was not found in SARS-CoV-2 spike glycoproteins.

A proper methodology was followed during the in silico designing of a multi-epitope vaccine candidate. The potential epitopes were screened using molecular docking and were selected based on the lowest binding affinity. The vaccine was constructed using these epitopes only for better binding results. This vaccine candidate shows effective binding with the HLA-B*15:03 subtype that is effective in eliciting an immune response against SARS-CoV-2. The in silico immune simulation of the designed vaccine showed a significant rise in TH cells after the first injection and reached its maximum level after the second injection accompanied by the formation of TH memory cells. Further, the total number of CTL and DCs increased over three months. IgM levels reached a high value after the second injection followed by a rise in IgG levels. Furthermore, IL-2 reached its maximum value at day 7 and then attained even higher levels after the second injection. These results show that the designed multi-epitope vaccine candidate is capable of inducing a humoral immune response against covid19. However, it still requires the experimental results and clinical trials to be more assured about the same. Additionally, this designed vaccine candidate could be beneficial in providing cross-protection against covid19 and the other similar viral diseases.

## Methods

### Phylogenetic analysis

Protein sequences of SARS-CoV-2 spike glycoprotein were retrieved from NCBI (www.ncbi.nih.nlm.gov) (Table 3). A phylogenetic tree of spike glycoprotein sequences of SARS-CoV-2 was constructed using IQTREE (L. T. Nguyen, Schmidt, Von Haeseler, & Minh, 2015). SARS-CoV spike glycoprotein sequences were used as the outgroup and were downloaded from NCBI (www.ncbi.nih.nlm.gov). The best model used was VT+G4, namely, the ‘Variable Time’ matrix (Müller & Vingron, 2001) with Gamma distribution (under four rate categories) as predicted by ModelFinder (Kalyaanamoorthy, Minh, Wong, Von Haeseler, & Jermiin, 2017). Some of the outliner (low scoring) sequences were removed according to the phylogenetic tree.

**Table 3.** Sequence details of spike glycoprotein sequences of SARS-CoV-2 that were obtained from phylogenetic analysis and were utilized in further analyses.

| Sr.no | Sequence ID/<br>PDB ID | Sequence Name | Chain | Sequence Length |
| --- | --- | --- | --- | --- |
| 1 | QIC53213.1 | spike_glycoprotein_SARS_corona_virus_2 | - | 1274 |
| 2 | QHR63290.2 | spike_glycoprotein_SARS_corona_virus_2 | - | 1274 |
| 3 | QHR63280.2 | spike_glycoprotein_SARS_corona_virus_2 | - | 1274 |
| 4 | QHR63270.2 | spike_glycoprotein_SARS_corona_virus_2 | - | 1274 |
| 5 | QHR63260.2 | spike_glycoprotein_SARS_corona_virus_2 | - | 1274 |
| 6 | QHR63250.2 | spike_glycoprotein_SARS_corona_virus_2 | - | 1274 |
| 7 | 6VYB | SARS-CoV-2_spike_glycoprotein | C | 1282 |
| 8 | 6VYB | SARS-CoV-2_spike_glycoprotein | B | 1282 |
| 9 | 6VYB | SARS-CoV-2_spike_glycoprotein | A | 1282 |
| 10 | 6VXX | SARS-CoV-2_spike_glycoprotein | C | 1282 |
| 11 | 6VXX | SARS-CoV-2_spike_glycoprotein | B | 1282 |
| 12 | 6VXX | SARS-CoV-2_spike_glycoprotein | A | 1282 |
| 13 | 6VSB | SARS-CoV-2_spike_glycoprotein | C | 1289 |
| 14 | 6VSB | SARS-CoV-2_spike_glycoprotein | B | 1289 |
| 15 | 6VSB | SARS-CoV-2_spike_glycoprotein | A | 1289 |
| 16 | 6LZG | SARS-CoV-2_Spike_receptor-binding_domain | B | 210 |
| 17 | 6W41 | Spike_glycoprotein_receptor_binding_domain | C | 232 |
| 18 | QIQ08769.1 | Surface_glycoprotein_partial_SARS_coronavirus | - | 53 |
| 19 | QIQ08768.1 | Surface_glycoprotein_partial_SARS_coronavirus2 | - | 53 |
| 20 | QII57161.1 | S_protein_SARS_coronavirus_2 | - | 1274 |
| 21 | QIA20044.1 | Surface_glycoprotein_SARS_coronavirus2 | - | 1274 |
| 22 | YP_009724390.1 | Surface_glycoprotein_SARS_coronavirus2 | - | 1274 |
| 23 | 6M0J | SARS-CoV-2_receptor-binding_domain | E | 230 |

#### Selection Analysis

The evolutionary analysis of SARS-CoV-2 spike glycoprotein sequences and GISAID sequences was carried out at the Datamonkey server (https://www.datamonkey.org) of HYPHY program (Pond & Muse, 2005). BUSTED (Branch-site Unrestricted Statistical Test for Episodic Diversification) (Murrell et al., 2015), FUBAR (Fast Unconstrained Bayesian Approximation) (Murrell et al., 2013) and MEME (Mixed Effects Maximum Likelihood) (Murrell et al., 2012) methods were used to detect episodic and pervasive selection pressure at each site respectively.

### In silico design of multi-epitope vaccine

#### Antigenic sequence identification

The antigenic sequences among the selected sequences were identified using the VaxiJen server (http://www.ddg-pharmfac.net/vaxijen/VaxiJen/VaxiJen.html). The threshold was set to the default value of 0.5. VaxiJen predicts protective antigens with accuracy ranging between 79% to 89% as tested on three different datasets (Doytchinova & Flower, 2007).

#### Epitope prediction and selection

B-cell epitopes were predicted using SVMTrip (Yao, Zhang, Liang, & Zhang, 2012). T-cell epitopes were predicted using Tepitool (Paul, Sidney, Sette, & Peters, 2016) of IEDB (Vita et al., 2019) with IC-50 value more than 500 nM for MHC-I and MHC-II. CTLs were predicted using CTLPred (Bhasin & Raghava, 2004). The predicted epitopes were subjected to immunogenicity analysis using IEDB (Vita et al., 2019). The predicted potential immunogens were analyzed for toxicity using ToxinPred (Gupta et al., 2013).

#### Structure prediction and validation of HLA-B*15:03 subtype

Homology modeling was performed to obtain the structures of HLA-B*15:03 subtype and selected epitopes using the Swissmodel (Waterhouse et al., 2018). The predicted structures of HLA-B*15:03 was further validated by creating the Ramachandran Plot using PROCHECK (Laskowski, MacArthur, Moss, & Thornton, 1993).

#### Selection of potential epitopes for multi-epitope vaccine

The selected epitopes with potential immunogenicity were docked with HLA-B*15:03 subtype to filter out epitopes. The docking was carried out using Autodock Vina (Trott & Olson, 2009). Binding pocket and residues in HLA-B*15:03 were identified using the CASTp 3.0 server (Tian, Chen, Lei, Zhao, & Liang, 2018). The binding x, y, and z coordinates in Autodock Vina were defined as 5.103, 13.862, and 146.304 respectively.

#### Construction of multi-epitope vaccine

Top 5 epitopes from each B-cell, T-cell, and CTL epitopes showing the lowest binding affinities towards HLA-B*15:03 were selected from the bound epitopes. The finally selected epitopes were joined together using GPGPG and AAY linkers. An adjuvant (Q0VDF9) was added at the N-terminal of the multi-epitope vaccine using the EAAAK linker to increase its immunogenicity. The adjuvant sequence was downloaded from the Uniprot database (https://www.uniprot.org).

#### Secondary and tertiary structure prediction of multi-epitope vaccine

Secondary structure was predicted by the PSIPRED server (http://globin.bio.warwick.ac.uk/psipred/) and RaptorX (http://raptorx.uchicago.edu/StructurePropertyPred/predict/). The PSIPRED first identifies homologs using PSI-BLAST and then employs feed-forward neural networks to predict the secondary structure (McGuffin, Bryson, & Jones, 2000). RaptorX predicts secondary structure without using any template. It incorporates deep convolutional neural fields to predict secondary structure, solvent accessibility, and disordered regions (S. Wang, Li, Liu, & Xu, 2016). The ab-initio tertiary structure of designed multi-epitope was predicted using I-Tasser (Yang & Zhang, 2015) server (https://zhanglab.ccmb.med.umich.edu/I-TASSER/). The I-Tasser (Iterative threading assembly refinement) predicts accurate structures by first identifying templates from PDB using multiple threading and identifies functions through the protein functions database. It has been ranked as the best server for protein structure and function prediction in CASP experiments (Shey et al., 2019). The predicted tertiary structure of the multi-epitope vaccine was validated using PROCHECK (Laskowski et al., 1993).

#### Physicochemical properties prediction of multi-epitope vaccine

The allergenicity of the designed vaccine candidate was predicted using Allertop v. 2.0 (http://www.ddg-pharmfac.net/AllerTOP) and AllergenFP v. 1.0 (http://ddg-pharmfac.net/AllergenFP). Allertop predicts allergenicity by incorporating the auto cross-variance (ACC) protein sequence mining method (Dimitrov, Bangov, Flower, & Doytchinova, 2014). AllergenFP calculates the Tanimoto coefficient to rate all protein pairs. Protein from the pair with the highest Tanimoto coefficient is classified as allergen or non-allergen (Dimitrov, Naneva, Doytchinova, & Bangov, 2014).

Physicochemical properties of designed candidate vaccine were predicted using Protparam (https://web.expasy.org/protparam/) (Gasteiger et al., 2005). Properties including theoretical PI, molecular weight, estimated half-life, instability index, and Grand Average of Hydropathicity (GRAVY) were estimated.

#### Molecular Docking of multi-epitope vaccine with HLA-B*15:03 subtype

Active residues of the multi-epitope vaccine were identified using the CASTp server (http://sts.bioe.uic.edu/castp/) (Tian et al., 2018). The multi-epitope vaccine was docked with HLA-B*15:03 using HADDOCK 2.4 webserver (https://bianca.science.uu.nl/haddock2.4/). HADDOCK (High Ambiguity Driven protein-protein Docking) is an integrated platform for flexible docking and performing short molecular dynamics simulations of molecular complexes (Van Zundert et al., 2016). The docking of multi-epitope and HLA-B*15:03 was performed by defining identified active residues. The active residues of HLA-B allele subtype predicted structure identified from CPORT server were *Gly80, Pro81, Glu82, Tyr83, Trp84, Asp85, Arg86, Glu87, Thr88, Gln89, Ile90, Ser91, Lys92, Thr93, Asn94, Thr95, Gln96, Thr97, Tyr98, Ala174, Arg175, Glu176, Ala177, Glu178, Gln179, Leu180, Arg181, Ala182, Tyr183, Leu184, Glu185, Gly186, Leu187, Cys188, Val189, Glu190, Trp191, Leu192, Arg193, Arg194, Tyr195, Leu196, Glu197, Asn198*, and *Gly199*. Active residues for multi-epitope vaccine identified using CPORT were *Tyr18, Phe281, Asp292, Ile377, Gly380, Leu385, Ser389, Leu390, Gly538, Gly542, Ala545, Glu546, Leu549, Ala550, Pro553, Gly554, Pro555, Asn559, Val560, Phe561, Ala562, Pro564, Gly566, Pro567, Gly568, Pro569, Gly570, Val571, Val572, Val573, Ala576, Asn594, His597, Ala599, Ser601, Phe604, Ala635, Cys639, Pro640, Phe641, Glu643, Val644, Phe645, Ala646, Ala647, Tyr648, Phe649, Gly650, Asp651, Asp652, Thr653, Val654, Ile655, Glu656, Val657, Ala658, Ala659, Tyr660, Phe661, Ser662, Tyr663, Phe664, Ala665, Val666, His667, Phe668, Ile669, Ala673, Val680, Val681, Ala682, Ala683, Tyr684, Lys685, Arg688, Leu691*, and *Ala692*. The passive residues were set to be detected in surrounding surface. An MD simulation of 1000 steps of energy minimization, 100 steps for heating phase, 1250 steps at 300 K, and 500 steps of cooling phase was performed.

#### In silico immune simulation

The immunogenicity and immune response of the designed multi-epitope vaccine was determined using the C-ImmSim server (http://150.146.2.1/C-IMMSIM/index.php?page=1). C-ImmSim is an agent-based simulator that uses systems biology and data-driven prediction methods to predict immune response to viruses, bacteria, or a vaccine (Rapin, Lund, Bernaschi, & Castiglione, 2010). The following most common alleles in Caucasian, Asian and Afro Brazilian were chosen HLA-A*02, 24, HLA-A*02, 03, DRB1*11, 13, DRB1*13, 11, 03 along with HLA-B*15:03 since it was shown to have great efficiency in presenting conserved SARS-CoV-2 that are shared between common human CoVs (A. Nguyen et al., 2020). The rest of the parameters were set as the following default values. Random Seed (12345) was used, simulation volume was set to 10, and simulation steps were set to 270 (= 90 days) with two injections the first at the 1-time step which equals the first eight hours of simulation, and the second injection at 63-time steps which is equal to three weeks. The rest of the parameters were left with default values. No specific antigen sequence was added to the inputs.

## Declaration

The authors declare that there is no conflict of interest whatsoever.

